# Trick or Treat? Pollinator attraction in *Vanilla pompona* (Orchidaceae)

**DOI:** 10.1101/2021.08.30.458254

**Authors:** Charlotte Watteyn, Daniela Scaccabarozzi, Bart Muys, Nele Van Der Schueren, Koenraad Van Meerbeek, Maria F. Guizar Amador, James D. Ackerman, Marco V. Cedeño Fonseca, Isler F. Chinchilla Alvarado, Bert Reubens, Ruthmery Pillco Huarcaya, Salvatore Cozzolino, Adam P. Karremans

## Abstract

Natural pollination of species belonging to the pantropical orchid genus *Vanilla* remains poorly understood. Based on sporadic records, euglossine bees have been observed visiting flowers of Neotropical *Vanilla* species. Our research aimed at better understanding the pollinator attraction mechanism of the Neotropical species *Vanilla pompona*, a crop wild relative with valuable traits for vanilla crop improvement programs. Using video footage, we identified floral visitors and examined their behavior. The flowers of *V. pompona* attracted *Eulaema cingulata* males, which distinctively displayed two behaviors: floral scent collection and nectar search; with the latter leading to pollen removal. Morphological measurements of floral and visitor traits showed that other *Eulaema* species may also act as potential pollinators. Additionally, we recorded natural fruit set in three populations and over a period of two years, tested for nectar presence and analyzed floral fragrances through gas chromatography - mass spectrometry. We observed a low natural fruit set (2.42%) and did not detect nectar. Twenty floral volatile compounds were identified, with the dominant compound trans-carvone oxide previously found to attract *Eulaema cingulata* males. We hypothesize a dual attraction of *Eulaema cingulata* males to *V. pompona* flowers, based on floral fragrance reward as the primary long-distance attraction, and food deception for successful pollen removal. Further research confirming this hypothesis is recommended to develop appropriate conservation policies for *Vanilla* crop wild relatives, which are the primary reserves of this crop’s genetic variation.

## 1. INTRODUCTION

Plant-pollinator interactions are typically mutualistic, with a reward offered by the plant in exchange for a pollination service (Wright & Schiestl, 2009). Within the orchid family (Orchidaceae), approximately two-thirds of all species reward their pollinators with nectar, floral fragrances, oils or food hairs (van der Pijl & Dodson, 1966; Nilsson, 1992; Williams, 1982; Shrestha et al., 2020), while the remaining are believed to be deceptive (Dafni, 1984; Ackerman, 1986). Among the floral fragrance-rewarding species, odor-seeking male orchid bees (Apidae: Euglossini) have been reported to be responsible for the pollination of hundreds of Neotropical orchid species (Dodson et al., 1969; Williams & Whitten, 1983). These males gather volatile compounds from floral or other sources to be used as courtship attractants, a phenomenon known as the “Perfume Flower Pollination Syndrome” (PFPS) (Dodson, 1975; Kimsey, 1984; Schemske & Lande, 1984; Eltz et al., 2003, 2005; Cappellari Rabeling & Harter-Marques, 2010).

The pantropical orchid genus *Vanilla* Plum. ex Mill. (1754) consists of 118 species (Karremans et al., 2020). Only those native to the Neotropics and belonging to the section *Xanata* produce aromatic pods or beans, such as *Vanilla planifolia* Andrews and its crop wild relatives (CWRs). Commercial vanilla plants are almost exclusively hand-pollinated, leading to satisfactory bean yields (Havkin-Frenkel & Belanger, 2018; Hu et al., 2019). Vector-mediated natural pollination, however, generally results in cross-pollination and increases genetic variability. The latter augments survival probabilities in changing environments and contributes to the development of potentially interesting crop traits related to, amongst others, pest resistance and bean quality. Studies on natural vanilla pollination are scarce but crucial to safeguard the long-term survival of this crop and its CWRs, which are the primary reserve of a crop’s genetic variation and a prerequisite for crop improvement programs (Castañeda-Álvarez et al., 2016; Flanagan & Mosquera-Espinosa, 2016).

The intrageneric diversity in floral morphology suggests the evolution of different pollination mechanisms (Cameron & Soto Arenas, 2003; Soto Arenas & Dressler, 2010). Even though spontaneous autogamy has been reported (Bateman et al., 2004; Van Dam et al., 2010; Pansarin, 2016), most *Vanilla* species are suspected to depend on animal vectors for pollination (Pansarin et al., 2014; Gigant et al., 2016; Chaipanich et al., 2020). Members of the Euglossini tribe are suggested as pollinators of the Neotropical species (Ackerman, 1983a; Lubinsky et al., 2006; Householder et al., 2010; Soto Arenas & Dressler, 2010; Pansarin & Pansarin, 2014; Anjos et al., 2017), but the specific mechanisms are still poorly understood. Our study aimed at better understanding the pollinator attraction mechanism and reproductive biology of the Neotropical species *Vanilla pompona* Schiede (*Vanilla* sect. *Xanata*), by a) identifying its floral visitors and comparing their behaviors, b) confirming its pollinator species and quantifying natural fruit set, c) identifying floral rewards, and d) predicting an optimal morphological fit between flower and pollinator to determine other potential pollinator species.

## 2. MATERIALS AND METHODS

### 2.1. Study area and species

The study took place in Costa Rica and Peru, in respectively the Osa Peninsula and the Madre de Dios region (Appendix A - Figure S1a-b). Both regions are recognized biological hotspots (Kohlmann et al., 2010) with a tropical wet climate (Atrium Biodiversity Information System, 2011; Holdridge, 1967; Kappelle, 2003) and host several *Vanilla* species (Householder et al., 2010; Janovec et al., 2013; Karremans et al., 2020; Watteyn et al., 2020).

*Vanilla pompona*, listed by the IUCN as Endangered (Herrera-Cabrera, 2020), has a distribution from Mexico to Brazil and Peru (Soto Arenas & Dressler, 2010). Flowers consist of three sepals, two lateral petals, and one large trumpet-shaped petal known as the lip or labellum. The latter is partially fused to the column, which bears a single ventral stigma, separated by the rostellum from the anther holding two pollinia-like pollen masses, that are deposited as smears. Each flower lasts for one day and starts to wither after midday.

### 2.3. Floral visitors and their behavior

Floral visitors were observed in the three Costa Rican and Peruvian study sites (hereafter *V. pompona* populations) during the flowering period of January-February 2019 (120 h of observation) and September 2018 (64 h of observation), respectively. We checked for newly-opened flowers and observed them from 0600 h to 1400 h, corresponding to the time of anthesis until the flowers start to wither. Cameras (CANON SX60 HS, EOS M) were located at a minimum distance of three meters from the flower to avoid disturbance. Videos were recorded upon visitor approaches (Costa Rica) or with tele macro mode activated (Peru). We screened the video footage to identify floral visitors, record their visiting time, and examine their behavioral patterns. Subsequently, we assigned them to five behavioral categories, following the behavioral hierarchy of bee attraction: (a) approaching, (b) landing on tepals or labellum, (c) landing on labellum, (d) entering labellum, and (e) successfully removing pollen. To model the effect of floral visitor genus and time of the day on this ordinal bee behavior classification, we used a series of mixed logistic regression models, and sequentially compared each pair of consecutive behavior categories (Appendix B) (Zuur et al., 2009).

### 2.4. Pollinator identification and pollination success

The floral visitors that removed pollen were identified at species level and determined by sex. Additionally, bees carrying pollen were caught during the flowering period in Costa Rica and Peru, using baits shown to be attractive compounds: eugenol, 1,8-cineole, and methyl salicylate (Dressler, 1981; Ackerman, 1989; Roubik & Hanson, 2004). The identity of the pollen was confirmed by morphological comparison with *V. pompona* pollen using high microscope magnification. Bees were identified using the key in Roubik and Hanson (2004). To test for autogamy, ten inflorescences from two Peruvian populations were bagged the day before anthesis with an insect proof nylon bag using a ground-based tool (Scaccabarozzi et al., 2020). Fruit set within the Costa Rican populations was quantified during two flowering periods (2019 and 2020). We counted the number of inflorescences per population, flowers per inflorescence, and developed fruits per inflorescence. The calculated fruit set was then compared among the three populations using the non-parametric Kruskal-Wallis test and Dunn’s post-hoc tests with multiple comparison correction (Benjamini-Hochberg adjustment to control the False Discovery Rate) and the two flowering periods using the non-parametric Mann-Whitney test.

### 2.5. Floral rewards

To determine if pollination is based on a nectar reward, we bagged flowers the day before anthesis (Corbet, 2003) and visually inspected 20 flowers from the Costa Rican populations for secretory glands under a stereoscopic microscope. Attempts to collect nectar using microcapillary tubes (2 μL) were undertaken in 10 of these flowers. Since no secreted fluids were observed or collected, we applied the rinsing method for small nectar volumes (Morrant et al., 2009; Power et al., 2017) in eight additional flowers. The rinsed water was analyzed using High-Performance Liquid Chromatography (HPLC) (Appendix C).

Floral fragrances of 12 flowers were collected by static headspace sampling (Tholl et al., 2006) during the 2020 flowering period in two Costa Rican populations. The sampling was conducted between 7:00 am and 11:00 am, corresponding to the activity peak of potential pollinators (Dodson et al., 1969; Ackerman, 1983b; Armbruster et al., 2000; Gostinski et al., 2016), thus expected floral scent productivity peaks (Whitten, 1985; Hills & Williams, 1990). Control samples were collected simultaneously from bags filled with ambient air. The scent traps were analyzed by gas chromatography-mass spectrometry (GC-MS). We used the same methodology for data analysis and processing, and compound identification and attractiveness as described in Hetherington-Rauth & Ramirez (2016) (Appendix C). The compound peak areas of the final set of floral volatiles were converted to relative frequencies and averaged per compound among the samples to reveal potential patterns within the floral bouquet.

### 2.6. Flower and floral visitor morphology

We measured and compared pollinator dimensions with floral traits to assess the pollinator morphological-fit to flower structure. Bee traps were located in the vicinity of the Peruvian and Costa Rican populations. The same methodology was used as described in Section 2.4. The bees’ total length, mesosoma height and diameter, and extended tongue length were measured using a caliper. Next, 32 *V. pompona* flowers were measured across the Costa Rican populations. Measured traits were: height (H1), width (W) and length (L1) of the labellar tube, height of the labellar tube below the anther (H2), thickness of the column at height of the anther (T_Co), height of the labellar callus (T_Ca), height of the labellar keels (T_La), and length of the pedicel (L2) (Figure 1a). Further derived variables were estimated to assess the size of the labellar tube: “An”, labellar tube entrance measured at the apex of the anther (H2 - T_La) and “Cn”, labellar tube measured at the height of the callus, below the rostellar flap (H2 - T_Ca).

**Figure 1.**
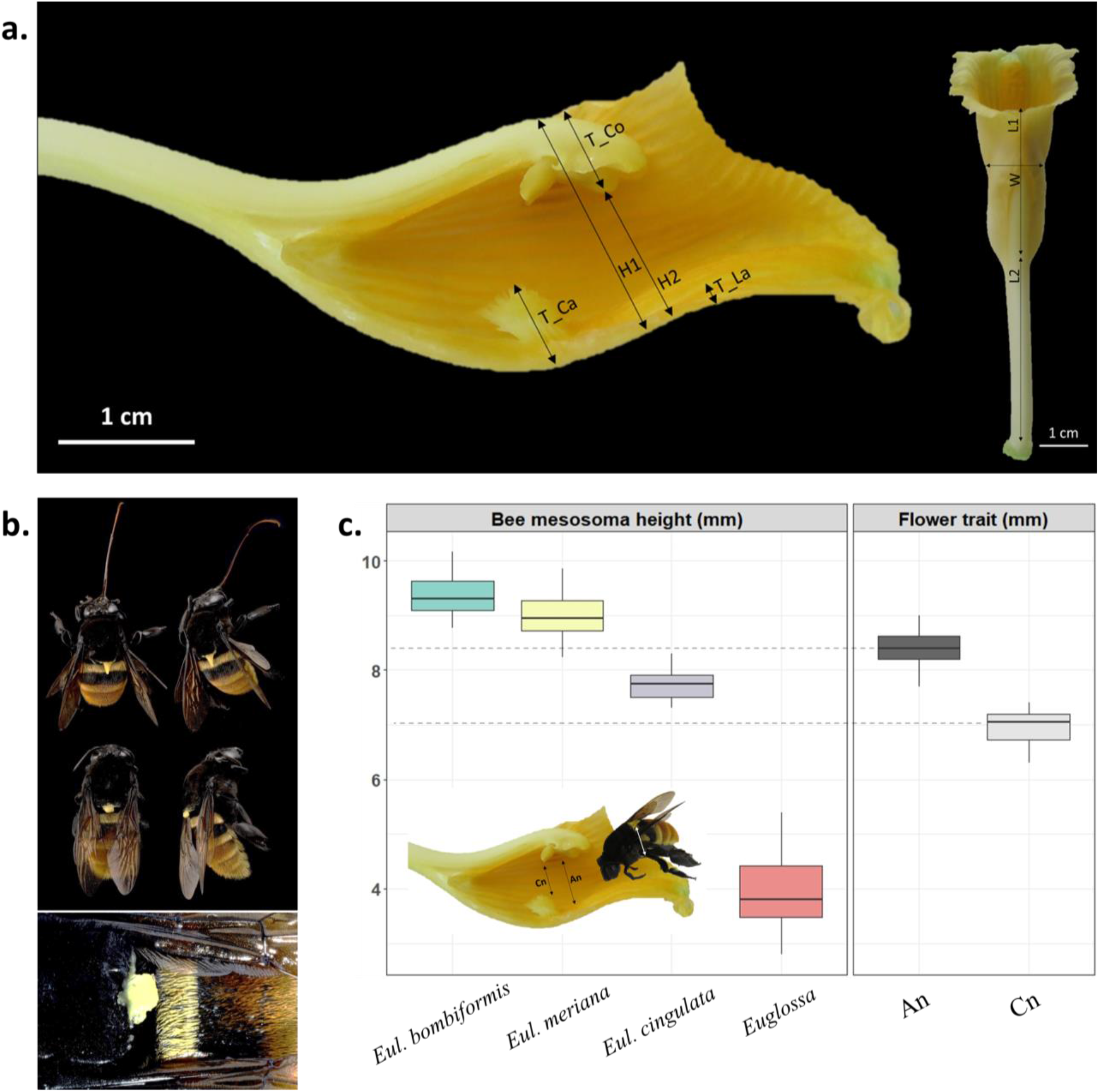
**(a)** Measured *V. pompona* floral traits: height (H1), width (W) and length (L1) of the labellar tube, height of the labellar tube below the anther (H2), thickness of the column at the height of the anther (T_Co), height of the labellar callus (T_Ca), height of the labellar keels (T_La), and length of the pedicel (L2). **(b)** Caught male *Eulaema cingulata* bees with attached pollen of *V. pompona* and extended tongue (top), and attached *V. pompona* pollen in close-up (bottom). **(c)** Comparison of the mesosoma heights (mm) of the bee species *Eulaema bombiformis*, *Eulaema meriana*, *Eulaema cingulata* and *Euglossa* spp. with the flower morphological traits “An” (mm) and “Cn” (mm) of *V. pompona*. Photographs by AK and CW (vouchers at Jardín Botánico Lankester and Museo de Entomología Klaus Raven Büller).

## 3. RESULTS

### 3.1. Floral visitors and their behavior

Of the 123 recorded floral visitors, 82.1% were Euglossini, belonging to the genera *Eulaema* (n = 73) and *Euglossa* (n = 28), and 17.9% were Trigonini of the genus *Trigona* (n = 22). The behavioral comparison analysis showed that most bees approaching the flowers subsequently landed on them (Appendix B - Table S1). Closer towards the end of the morning, the chance that a bee lands on a flower decreases significantly (Appendix B - Figure S2). Approaching *Euglossa* or *Eulaema* are predicted to land more on the tepals than on the labellum, where they show a probing behavior of brushing the flower structure with their foretarsi (Appendix B - Table S2). The chance that an approaching *Trigona* lands on the labellum edge or enters the labellar tube is higher compared to *Euglossa* or *Eulaema*, and all *Trigona* display a probing behavior of touching the flower surface with their proboscis or antennae. *Eulaema* or *Euglossa* that enter the labellum, visit the flower for less than five seconds and remain inside for about two seconds. As they exit, an extended tongue was observed. These rapid visits suggest unsuccessful attempts of nectar foraging. Despite the observation of three different bee genera in both Costa Rica and Peru, only individuals belonging to the genus *Eulaema* (n = 5) successfully removed pollen when entering the labellum (Appendix B - Figure S3). Overall, landing bees display two main reward-approach behaviors on different flower parts: probing (Video S1) or attempts for nectar search (Video S2).

### 3.2. Pollinator identification and pollination success

The five bees observed removing the pollen of *V. pompona* flowers were identified as male *Eulaema cingulata*. Furthermore, using chemical baits, we caught five male *Eulaema cingulata* that carried *V. pompona* pollen masses attached to their scutellum (Figure 1b). Fruit set at inflorescence level was significantly different among the populations (*H*(2) = 7.72, *P* = 0.02), with Piro-Filanance (4.75% ± 9.71%) having a higher fruit set compared to Miramar (1.26% ± 4.78%; *P* = 0.01) and Piro-Rio (1.24% ± 4.57%; *P* = 0.02). No difference was found between the two years (*H*(1) = 0.42, *P* = 0.52). The autogamy test resulted in zero fruit development.

### 3.3. Floral rewards

No nectaries were observed after visually inspecting 20 *V. pompona* flowers and no nectar could be collected via microcapillary tubes. The rinsing method followed by HPLC analysis detected total sugars (glucose, fructose, sucrose) in trace (< 0.03 mg) amounts (four samples) or non-quantifiable amounts (five samples) (Appendix C - Figure S4). We detected a total of 20 floral fragrance compounds emitted by *V. pompona* flowers (Table 1). One of them could not be identified. The most representative compounds of the floral bouquet were trans-carvone oxide (35.70 ± 14.85%), limonene (17.45 ± 6.00%), limonene oxide, trans (16.72 ± 6.87%) and limonene oxide, cis (8.97 ± 6.80%), followed by terpinolene (6.94 ± 3.60%), carvone (4.42 ± 1.50%), myrcene (3.34 ± 2.36%), 1-R-alpha-pinene (2.61 ± 2.30%) and (-)-beta-pinene (1.33 ± 0.93%) (Appendix C - Figure S5). The other compounds were present in low amounts (mean relative peak area < 1%).

**Table 1.**
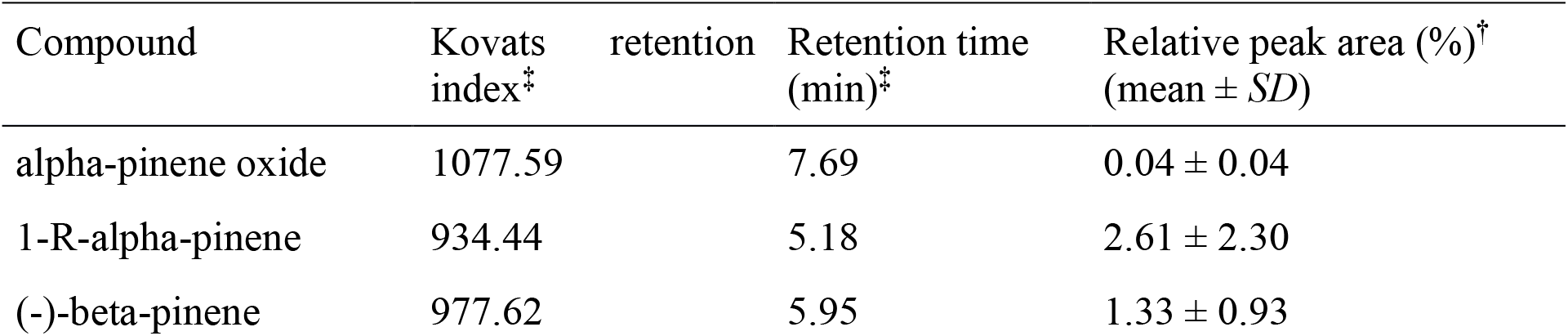
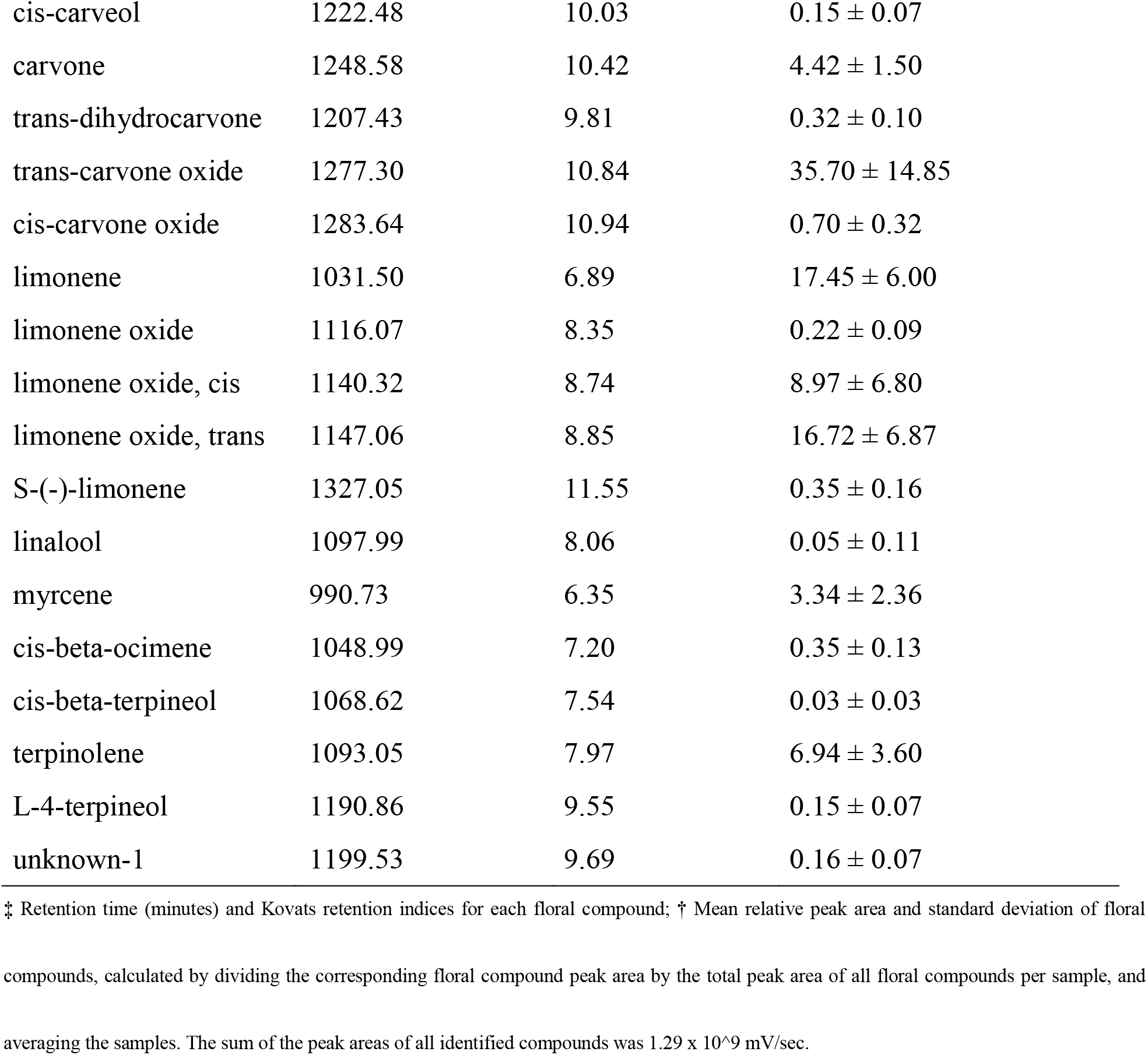
Identified floral volatile compounds, Kovats retention indices, retention times (min), and mean relative peak areas (%) calculated from the total peak area in the GC-MC chromatogram from *Vanilla pompona* flowers (*N* =12). *SD* = standard deviation.

### 3.4. Flower and floral visitor morphology

A total of 83 *Eulaema* (3 species), 690 *Euglossa* (20 species), and 32 *V. pompona* flowers were measured. The trait “An” (8.40 ± 0.35 mm) and “Cn” (6.95 ± 0.36 mm) showed the best fit with the height of the pollinator species *Eulaema cingulata* (7.73 ± 0.31 mm) (Figure 1c). In fact, *Eulaema cingulata* fits exactly between these two traits. The *Euglossa* species (3.95 ± 0.58 mm) were too small to fall into the An-Cn gap, whereas the other *Eulaema* species, *Eulaema bombiformis* (9.39 ± 0.39 mm) and *Eulaema meriana* (8.98 ± 0.44 mm), were too large.

## 4. DISCUSSION

*Vanilla pompona* flowers are visited by bees of the genera *Eulaema*, *Euglossa* and *Trigona*. As confirmed by both the autogamy test and video footage, sexual reproduction is vector-mediated. Only male *Eulaema cingulata* were observed removing pollen and caught with *V. pompona* pollen masses. They are therefore suspected to act as effective pollinators in both Costa Rica and Peru. Yet, it is not possible to exclude females nor other species as potential pollinators.

Bee behavioral patterns were assessed and compared. Towards the end of the morning, a decrease in number of landing bees suggests that flowers become less attractive due to senescence close to midday or after previous floral visits, as reported earlier for several plant species (van Doorn, 1997). *Trigona* landed almost exclusively on the labellum and exhibited a probing behavior indicative of foraging (Armbruster, 1984; Slaa et al., 1998; Nieh, 2004; Lichtenberg et al., 2010; McCabe & Farina, 2010; Grajales-Conesa et al., 2012). *Euglossa* and *Eulaema* were observed more frequently on the tepals, repeatedly brushing with their foretarsi, then hovering around while transferring compounds to their hind tibia, indicating fragrance collection (Dressler, 1968; Whitten et al., 1989; Eltz et al., 2003). Some *Euglossa* and *Eulaema* did not collect fragrances but entered directly into the labellum. These visits were brief, after which they quickly exited the flower showing an extended tongue, suggesting nectar search. Only *Eulaema cingulata* successfully removed the pollen after searching for nectar in the labellum.

### 4.1. The dual pollinator attraction hypothesis

The floral volatile compound composition was quantified for the first time in *V. pompona*. Trans-carvone oxide, limonene and limonene oxide together represented 78.84% of the floral perfume. Previous field bioassays have demonstrated that the compound trans-carvone oxide attracts *Eulaema cingulata* and other *Eulaema* species, that collect and store it in their hind legs (Whitten et al., 1986, Etlz et al., 2006; Milet-Pinheiro & Gerlach, 2017). Flowers of *Catasetum* species, known to be pollinated through PFPS by different *Eulaema* species, also contain trans-carvone oxide (Whitten et al., 1986; Gerlach & Schill, 1991; Milet-Pinheiro & Gerlach; 2017). Hence, this compound most likely evolved in perfume-rewarding plant species as an attractant for *Eulaema* pollinator species (Whitten et al., 1986; Brandt et al., 2019). The specific role of the other identified compounds and their interaction with the *Eulaema* attractant trans-carvone oxide could be explored further since studies have demonstrated that appropriate compound blends filter out certain bee pollinator species (Williams & Dodson, 1972; Whitten et al., 1986).

For pollination to occur, bees need to actively crawl into the labellum. Curiously, none of the observed male *Eulaema cingulata* exhibiting fragrance collection behavior on the tepals subsequently moved into the labellum. We distinctly observed male *Eulaema cingulata* entering the labellum and exiting the flowers with extended tongues, indicative of nectar search. The identification of sugar traces suggests that *V. pompona* flowers deceive their pollinators, which is conform with our observations of scarce and speedy visits in the labellum and a low natural fruit set (2.42%) (Scopece et al., 2010; Scaccabarozzi et al., 2020). To tease apart whether the deception is based on resembling a general flower image or a specific model species (Johnson and Schiestl, 2016; Caballero-Villalobos et al., 2017), advertising signals shared by *V. pompona* and co-occurring *Eulaema cingulata*-pollinated species could be explored. Sugar traces most likely encourage the bees to crawl inside the labellum, but there exists quite some debate whether to decide if this is a food deceptive or semi-rewarding strategy. In case of a semi-rewarding strategy, however, we would expect revisits since traces do not satisfy the bees’ high energy demands. In accordance with Shrestha et al. (2020), we suggest that sugar traces are deceitful, but the nature of the deceit is more complex compared to complete nectar absence and definitely requires a more in-depth comprehension.

Combining our behavioral and phytochemical data, we hypothesize a dual pollinator attraction in *V. pompona* flowers, employing floral fragrance rewards as initial attraction and food deception to induce pollen removal and deposition. The volatile compounds, especially trans-carvone oxide, possibly provoke the primary long-distance attraction of male *Eulaema cingulata* to a flowering population. *V. pompona* flowers only last for one day and start to wither after midday. This restricted time lapse may have led to the development of floral fragrance rewards to boost pollinator attraction. Yet, the attempt for nectar search is essential for pollen removal and seems to occur via food deception as the short-distance attraction, taking place when these male bees, now present in the surroundings of the flowering *V. pompona*, are foraging for food resources. Studies involving flower manipulation experiments (i.e. removing fragrant compounds) or bee tagging could confirm our proposed hypothesis and examine if this mechanism has been evolved across other Euglossini-pollinated *Vanilla* species.

### 4.2. Morphological goodness-of-fit for pollen removal by *Eulaema*

We assessed the “goodness-of-fit” between flower and bee to ensure pollen removal and showed that the body height of the pollinator *Eulaema cingulata* perfectly fits in between the flower traits “An” and “Cn”. Given that vanilla flowers have a single aperture for entrance and exit, the callus serves as a positioning structure, bringing the bee close to the rostellum. This results in the latter being moved backwards, displacing the anther and releasing the pollen on the bee’s scutellum. It explains why the height of the labellar tube at the callus is slightly smaller than the bee height and demonstrates the key role of the labellar tube and the callus, both highly conserved features in *Vanilla* sect. *Xanata*. Since the chemical baits did probably not attract all available *Eulaema* species and the floral and bee morphological traits show intraspecific variation, more bee species could fall within the “An-Cn gap”. This is beneficial from a plant’s perspective since dependence on a single pollinator species might hazard a plant’s population survival, especially when subjected to environmental pressures (González-Varo et al., 2013; Traveset et al., 2017).

## 5. CONCLUSION

We hypothesize that *V. pompona* pollination by male *Eulaema cingulata* is based on floral fragrance attraction towards a flowering population and food deception to induce pollen removal and deposition. This proposed dual pollinator attraction mechanism can be explained by the need to (a) attract bees from long distances to flowering populations, and (b) lure bees into the labellum for successful pollination. Our study offers novel insights into plant-pollinator interactions within the *Vanilla* genus and could be corroborated by further research on the attraction of both sexes to *V. pompona* flowers, the presence of sugar traces as a semi-deceptive mechanism, the pollinator species diversity across *V. pompona*’s distribution within the Neotropics, and the effects of environmental changes on these interactions. Pollination studies are essential to define proper conservation policies, thereby preserving the natural habitat of both vanilla and its pollinators and assuring their long-term survival and agricultural application.

## ACKNOWLEDGEMENTS

CW, DS, BM, BR, SC and APK developed the experimental design of the research. CW, DS, NVDS, IC, MCF and APK collected the data. CW, DS, NVDS, KVM and APK analysed and interpreted the collected data. CW, DS and APK wrote the first draft of the paper. All other authors reviewed and approved the final paper. We would like to give a special thanks to José Pérez Fuentes, Jessica Fonseca Prendas, Marvin Lopez Morales, Andrea Aromatisi, Manuel Huinga Escalante, Luis Llerena and Osa Conservation for their help during the fieldwork, Mauricio Fernández to determine the sex of the *Eulaema* specimens, the Ramirez Lab to help us with analysing floral volatiles, the Insectopia lab and Clorinda Vergara Cobian for their help with bee identification, and the environmental associations Arbio Peru, Simbio Onlus, Camino Verde.

## DATA AVAILABILITY STATEMENT

Upon acceptance, the data that supports the findings of this study will be made openly available in the Dryad Digital Repository.

## SUPPORTING INFORMATION

Additional Supporting Information may be found online in the Supporting Information section.

## SUPPORTING INFORMATION

### Appendix A – Study area and study sites in Costa Rica and Peru

**Figure S1.** Map showing the study sites in (a) Osa Peninsula, Costa Rica (Miramar 8°26’20” N, 83°21’54” W, Piro 8°24’5” N, 83°20’36” W), and (b) Madre de Dios region, Peru (Baltimore 13°23’38” S, 69°44’59” W, Kapievi 13°1’17 S, 69°20’4” W, Belo Horizonte 12°28’38” S, 69°2’59” W).

### Appendix B – Mixed logistic regression models for ordinal bee behavior classification

**Table S1.** Output of Bayesian multilevel behavioral comparison models to sequentially compare behavioral categories between floral visitor genera and time of the day. The intercept term is the genus *Euglossa* of the categorical variable bee genus. LOOIC = leave-one-out cross-validation, WAIC = Widely Applicable Information Criteria, *SE* = standard error, CI = credibility interval.

**Table S2.** Percentage of landing bees per genus displaying a defined reward approach type on different parts of *Vanilla pompona* flowers (T = tepals; LE = labellum edge; LT = labellum tube), whereby “probing” refers to touching the flower surface with the proboscis and antennae and / or brushing with foretarsi, and “nectar search” refers to entering straight into the labellum chamber, and showing an extended tongue when entering or departing the flower. *N* = total number of landing bees per genus. *SD* = standard deviation.

**Table S3.** Mean fruit set (%) of *Vanilla pompona* populations at the three study sites in Costa Rica, measured during the flowering period of 2019 and 2020. *SD* = standard deviation.

**Figure S2.** Response curve of the behavior landing, shown over time and per bee genus; demonstrating the chance over time of the day that a bee individual belonging to a certain bee genus will land on a *Vanilla pompona* flower.

**Figure S3.** Percentage of floral visitors per bee genus that (1) approaches, i.e. shows interest but not necessarily lands, (2) lands, (3) probes, (4) enters into the labellum, (5) successfully removes the pollen of *V. pompona* flowers.

### Appendix C – Floral rewards

**Figure S4.** HPLC-chromatogram of a *V. pompona* sample extracted from the labellum via three successive rinses (0.5 mL each) of de-ionized UHPLC gradient water. The green line represents a standard mixture of three sugars (sucrose, glucose, fructose). The blue line shows the sugar content of the sample. The limits of detection (LOD) was determined by the signal to noise (S/N) ratio method and estimated as the minimum concentration providing a S/N ratio of 3:1.

**Figure S5.** Floral volatile compound composition of *Vanilla pompona* flowers, calculated by averaging the relative frequency of each individual compound among the twelve samples.

### Supporting Videos

**Video S1.** *Eulaema* and *Euglossa* demonstrating a probing behavior on *Vanilla pompona* flowers, i.e. collecting fragrances from the tepals (sepals and lateral petals).

**Video S2.** *Eulaema cingulata* entering the labellum of a *Vanilla pompona* flower as an attempt for nectar search, demonstrated by the extended tongue when entering and exiting the floral tube, and successfully removing the pollen in case of a morphological fit between bee and flower.

## REFERENCES

Ackerman, J. D. (1983a). Specificity and mutual dependency of the orchid-euglossine bee interaction. Biological Journal of the Linnean Society, 20(3), 301–314.

Ackerman, J. D. (1983b). Diversity and seasonality of male euglossine bees (Hymenoptera: Apidae) in Central Panama. Ecology, 64(2), 274–283.

Ackermann J. D. (1986). Mechanisms and evolution of food-deceptive pollination systems in orchids. Lindleyana, 1(2), 108–113.

Ackerman, J. D. (1989). Geographic and seasonal variation in fragrance choices and preferences of male euglossine bees. Biotropica, 21(4), 340–347.

Anjos, A. M., Barberena, F. F. V. A., & Pigozzo, C. M. (2017). Biologia reprodutiva de *Vanilla bahiana* Hoehne (Orchidaceae). Orquidário, 30(3/4), 67–79.

Armbruster, W. S. (1984). The role of resin in angiosperm pollination: ecological and chemical considerations. American Journal of Botany, 71(8), 1149–1160.

Armbruster, W. S., Fenster, C., & Dudash, M. (2000). Pollination “principles” revisited: specialization, pollination syndromes, and the evolution of flowers. The Scandanavian Association for Pollination Ecology, 39, 179–200.

Atrium Biodiversity Information System (2011). Retrieved from http://atrium.andesamazon.org

Bateman, R. M., Pridgeon, A. M., Cribb, P., Chase, M., & Rasmussen, F. N. (2004). Genera Orchidacearum Volume 3, Orchidoideae (Part 2), Vanilloideae. Kew Bulletin, 59(1). doi: 10.2307/4111088

Brandt, K., Dötterl, S., Fuchs, R., do Amaral Ferraz Navarro, D. M., Sobreira Machado, I. C., Dobler, D., Ayasse, M., & Milet-Pinheiro, P. (2019). Subtle chemical variations with strong ecological significance: stereoselective responses of male orchid bees to stereoisomers of carvone epoxide. Journal of Chemical Ecology, 45, 464–473.

Caballero-Villalobos, L., Silva-Arias, G. A., Buzatto, C. R., Nervo, M. H., & Singer, R. B. (2017). Generalized food-deceptive pollination in four *Cattleya* (Orchidaceae: Laeliinae) species from Southern Brazil. Flora, 234, 195–206.

Cameron, K. M., & Soto Arenas, M. A. (2003). Vanilloideae. In: Pridgeon AM, Cribb PJ, Chase MW, Rasmussen FN, eds. Genera Orchidacearum, vol 3, Orchidoideae, part 2, Vanilloideae, Oxford: Oxford University Press, 281–334.

Cappellari Rabeling, S., & Harter-Marques, B. (2010). First report of scent collection by male orchid bees (Hymenoptera: Apidae: Euglossini) from terrestrial mushrooms. Journal of the Kansas Entomological society, 83(3), 264–266.

Castañeda-Álvarez, N. P., Khoury, C. K., Achicanoy, H. A., Bernay, V., Dempewolf, H., Eastwood, R. J., Guarino, L., Harker, R. H., Jarvis, A., Maxted, N., Müller, J. V., Ramirez-Villegas, J., Sosa, C. C., Struik, P. C., Vincent, H., & Toll, J. (2016). Global conservation priorities for crop wild relatives. Nature Plants, 2(4). doi: 10.1038/nplants.2016.22

Chaipanich, V. V., Wanachantararak, P., & Hasin, S. (2020). Floral morphology and potential pollinator of *Vanilla siamensis* Rolfe ex Downie (Orchidaceae: Vanilloideae) in Thailand. Thailand Natural History Museum Journal, 14(1), 1–14.

Corbet, S. A. (2003). Nectar sugar content: estimating standing crop and secretion rate in the field. Apidologie, 34(1), 1–10.

Dafni, A. (1984). Mimicry and deception in pollination. Annual Review of Ecology and Systematics, 15(1), 259–278.

Dodson, C. H., Dressler, R. L., Hills, H. G., Adams, R. M., & Williams, N. H. (1969). Biologically active compounds in orchid fragrances. Science, 164 (3885), 1243–1249.

Dodson, C. H. (1975). Coevolution of orchids and bees. In: Gilbert, L. and Raven P. H., eds. Coevolution of animals and plants. Texas: University of Texas Press, 91–99.

van Doorn, W. G., & Stead, A. D. (1997). Abscission of flowers and floral parts. Journal of Experimental Botany, 48(4), 821–837.

Dressler, R. L. (1968). Pollination by euglossine bees. Evolution, 22(1), 202–210.

Dressler, R. (1981). The orchids - natural history and classification. Cambridge: Harvard University Press.

Eltz, T., Roubik, D. W., & Whitten, W. M. (2003). Fragrances, male display and mating behaviour of *Euglossa hemichlora*: a flight cage experiment. Physiological Entomology, 28(4), 251–260.

Eltz, T., Sager, A., & Lunau, K. (2005). Juggling with volatiles: exposure of perfumes by displaying male orchid bees. Journal of Comparative Physiology A, 191(7), 575–581.

Flanagan, N. S., & Mosquera-Espinosa, A. T. (2016). An integrated strategy for the conservation and sustainable use of native *Vanilla* species in Colombia. Lankesteriana, 16(2), 201–218.

Gerlach, G., & Schill, R. (1991). Composition of orchid scents attracting euglossine bees. Botanica Acta, 104(5), 379–384.

Gigant, R. L., De Bruyn, A., M’sa T, Viscardi, G., Gigord, L., Gauvin-Bialecki, A., Pailler, T., Humeau, L., Grisoni, M., & Besse, P. (2016). Combining pollination ecology and fine-scale spatial genetic structure analysis to reveal the reproductive strategy of an insular threatened orchid. South African Journal of Botany, 105, 25–35.

González-Varo, J. P., Biesmeijer, J. C., Bommarco, R., Potts, S. G., Schweiger, O., Smith H. G., Steffan-Dewenter, I., Szentgyorgi, H., Woyciechowki, M., & Vilà, M. (2013). Combined effects of global change pressures on animal-mediated pollination. Trends in Ecology and Evolution, 28(9), 524–530.

Gostinski, L. F., Carvalho, G. C. A., Rego, M. M. C., & Albuquerque, P. M. C. (2016). Species richness and activity patterns of bees (Hymenoptera: Apidae) in the restinga area of Lençóis Maranhenses National Park, Barreirinhas, Maranhão, Brazil. Revista Brasileira de Entomologia, 60(4), 319–327.

Grajales-Conesa, J., Meléndez Ramírez, V., Cruz-López, L., & Sánchez Guillén, D. (2012). Effect of Citrus floral extracts on the foraging behavior of the stingless bee *Scaptotrigona pectoralis* (Dalla Torre). Revista Brasileira de Entomologia, 56(1), 76–80.

Havkin-Frenkel, D., & Belanger, F. C. (2018). Handbook of Vanilla science and Technology, 2^nd^ edn. Ames: Blackwell Publishing Ltd.

Herrera-Cabrera, B. E., Hernández, M., Vega, M., & Wegier, A. (2020). The IUCN Red List of Threatened Species. Retrieved from https://www.iucnredlist.org/species/105878897/173977322.

Hetherington-Rauth, M. C., & Ramirez, S. R. (2016). Evolution and diversity of floral scent chemistry in the euglossine bee-pollinated orchid genus *Gongora*. Annals of Botany, 118(1), 135–148.

Hills, G. H., & Williams, N. H. (1990). Fragrance cycle of *Clowesia rosea*. Orquidea, 12, 119–132.

Holdridge, L. R. (1967). Life zone ecology, 1^st^ edn. San José: Tropical Science Center.

Householder, E., Janovec, J., Balarezo Mozambite, A., Huinga Maceda, J., Wells, J., Valega, R., (2010). Diversity, natural history, and conservation of *Vanilla* (orchidaceae) in amazonian wetlands of Madre de Dios, Peru. Journal of the Botanical Research Institute of Texas, 4(1), 227–243.

Hu, Y., Resende Jr., M. F. R., Bombarely, A., Brym, M., Bassil, E., & Chambers, A. H. (2019). Genomics-based diversity analysis of *Vanilla* species using a *Vanilla planifolia* draft genome and Genotyping-By-Sequencing. Scientific Reports, 9. doi: 10.1038/s41598-019-40144-1

Janovec, J., Householder, J., & Tobler, M. (2013). Evalucación de los actuales impactos y amenazas inminentes en aguajales y cochas de Madre de Dios, Perú. Lima: WWF.

Johnson, S. D., & Schiestl, F. P. (2016). Floral mimicry, 1^st^ edn. Oxford: Oxford University Press.

Kappelle, M. (2003). Ecosistemas del Area de Conservación Osa (ACOSA), 1st edn. Heredia: INBio

Karremans, A. P., Chinchilla, I. F., Rojas-Alvarado, G., Fonseca, M. C., Damian, A., Léotard, G. (2020). A reappraisal of Neotropical *Vanilla*. With a note on taxonomic inflation and the importance of alpha taxonomy in biological studies. Lankesteriana, 20(3), 395–497.

Kimsey, L. S. (1984). The behavioural and structural aspects of grooming and related activities in euglossine bees (Hymenoptera: Apidae). Journal of Zoology, 204, 541–550.

Kohlmann, B., Roderus, D., Elle, O., Solís, A., Soto, X., & Russo, R. (2010). Biodiversity conservation in Costa Rica: A correspondence analysis between identified biodiversity hotspots (Araceae, Arecaceae, Bromeliaceae, and Scarabaeinae) and conservation priority life zones. Revista Mexicana de Biodiversidad.

Lichtenberg, E. M., Imperatriz-Fonseca, V. L., & Nieh, J. C. (2010). Behavioral suites mediate group-level foraging dynamics in communities of tropical stingless bees. Insectes Sociaux, 57(1), 105–113.

Lubinsky, P., Van Dam, M., & Van Dam, A. (2006). Pollination of *Vanilla* and evolution in Orchidaceae. Orchids, 75(12), 926–929.

McCabe, S. I., & Farina, W. M. (2010). Olfactory learning in the stingless bee *Tetragonisca angustula* (Hymenoptera, Apidae, Meliponini). Journal of Comparative Physiology, 196(7), 481–490.

Milet-Pinheiro, P., & Gerlach, G. (2017). Biology of the Neotropical orchid genus *Catasetum*: a historical review on floral scent chemistry and pollinators. Perspectives in Plant Ecology Evolution and Systematics, 27, 23–34.

Morant, D. S., Schumann, R., & Petit, S. (2009). Field methods for samping and storing nectar from flowers with low nectar volumes. Annals of Botany, 103(3), 533–542.

Nieh, J. (2004). Recruitment communication in stingless bees (Hymenoptera, Apidae, Meliponini). Apidologie, 35(2), 159–182.

Nilsson, L. A. (1992). Orchid pollination biology. Trends in Ecology and Evolution, 7(8), 255–259.

Pansarin, E. R. & Pansarin, L. M. (2014). Floral biology of two Vanilloideae (Orchidaceae) primarily adapted to pollination by euglossine bees. Plant Biology, 16(6), 1104–1113.

Pansarin, E. R., Aguiar, J. M., & Pansarin, L. M. (2014). Floral biology and histochemical analysis of *Vanilla edwallii* Hoehne (Orchidaceae: Vanilloideae): an orchid pollinated by *Epicharis* (Apidae: Centridini). Plant Species Biology, 29(3), 242–252.

Pansarin, E. R. (2016). Recent advances on evolution of pollination systems and reproductive biology of Vanilloideae (Orchidaceae). Lankesteriana, 16(2), 255–267.

Power, E. F., Stabler, D., Borland, A. M., Barnes, J., & Wright, G. A. (2018). Analysis of nectar from low-volume flowers: A comparison of collection methods for free amino acids. Methods in ecology and evolution, 9(2), 734–743.

Roubik, D. W., & Hanson, P. E. (2004). Orchid bees of Tropical America: Biology and Field Guide, 1^st^ edn. Heredia: INBio.

Scaccabarozzi, D., Campbell, T., & Dods, K. (2020a). A simple and effective ground-based tool for sampling tree flowers at height for subsequent nectar extraction. Methods in Ecology and Evolution, 11(11), 1421–1426.

Scaccabarozzi, D., Galimberti, A., Dixon, K. W., & Cozzolino, S. (2020b). Rotating arrays of orchid flowers: a simple and effective method for studying pollination in food deceptive plants. Diversity, 12(8), 286.

Schemske, D., & Lande, R. (1984). Fragrance collection and territorial display by male orchid bees. Animal Behavior, 32(3), 264–266.

Scopece, G., Cozzolino, S., Johnson, S. D., & Schiestl, F. P. (2010). Pollination efficiency and the evolution of specialized deceptive pollination systems. The American Naturalist, 175(1), 98–105.

Shrestha, M., Dyer, A. G., Dorin, A., Ren, Z. X., & Burd, M. (2020). Rewardlessness in orchids: how frequent and how rewardless? Plant Biology, 22(4), 555–561.

Slaa, E. J., Cevaal, A., & Sommeijer, M. J. (1998). Flower constancy in *Trigona* stingless bees (Apidae, Meliponini) foraging on artificial flower patches: a comparative study. Journal of Apicultural Research, 37, 191–198.

Soto Arenas, M. A., & Dressler, R. L. (2010). A revision of the Mexican and Central American species of *Vanilla* plumier ex miller with a characterization of their its region of the nuclear ribosomal DNA. Lankesteriana, 9(3), 285–354.

Traveset, A., Tur, C., & Equíluz, V. M. (2017). Plant survival and keystone pollinator species in stochastic coexstinction models: role of intrinsic dependence on animal-pollination. Scientific Reports, 7(1), 6915.

Tholl, D., Boland, W., Hansel, A., Loreto, F., Röse, U. S. R., & Schnitzler, J. P. (2006). Practical approaches to plant volatiles analysis. The Plant Journal, 45(4), 540–560.

Van Dam, A. R., Householder, J. E., & Lubinsky, P. (2010). *Vanilla bicolor* Lindl. (Orchidaceae) from the Peruvian Amazon: Auto-fertilization in Vanilla and notes on floral phenology. Genetic Resources and Crop Evolution, 57(4), 473–480.

Van der Pijl, L., & Dodson, H. (1966). Orchid flowers: Their pollination and evolution, 1^st^ edn. Miami: University of Miami.

Watteyn, C., Fremout, T., Karremans, A. P., Pillco Huarcaya, R., Azofeifa Bolaños, J. B., Reubens, B., & Muys, B. (2020). Vanilla distribution modeling for conservation and sustainable cultivation in a joint land sparing/sharing concept. Ecosphere, 11(3). doi: 10.1002/ecs2.3056

Whitten, W. M. (1985). Variation in floral fragrances and pollinators in the Gongora quinquenervis complex (Orchidaceae) in Central Panama. PhD Thesis, Florida University, USA.

Whitten, W. M., Williams, N. H., Armbruster, W. S., Battiste, M. A, Strekowski, L., & Lindquist, N. (1986). Carvone oxide: an example of convergent evolution in euglossine pollinated plants. Systematic Botany, 11(1), 222–228.

Whitten, W. M., Young, A. M., & Williams, N. H. (1989). Function of glandular secretions in fragrance collection by male euglossine bees (Apidae: Euglossini). Journal of Chemical Ecology, 15(4), 1285–1295.

Williams, N. H., & Dodson, C. H. (1972). Selective attraction of male euglossine bees to orchid floral fragrances and its importance in long-distance pollen flow. Evolution, 26(1), 84–95.

Williams, P. H. (1982). The distribution and decline of British bumble bees (*Bombus* Latr.). Journal of Apicultural Research, 21(4), 236–245.

Williams, P. H., & Whitten, W. M. (1983). Orchid floral fragrances and male euglosssine bees: methods and advances in the last sesquidecade. Biological Bulletin, 164(3), 355–395.

Wright, G. A., & Schiestl, F. P. (2009). The evolution of floral scent: The influence of olfactory learning by insect pollinators on the honest signalling of floral rewards. Functional Ecology, 23(5), 841–851.

Zuur, A., Ieno, E. N., Walker, N., Saveliev, A. A, & Smith, G. M. (2009). Mixed effects models and extensions in ecology with R, 1st edn. New York: Springer.

