## Appendix A for "Trick or Treat? Pollinator attraction in *Vanilla pompona* (Orchidaceae)"

---

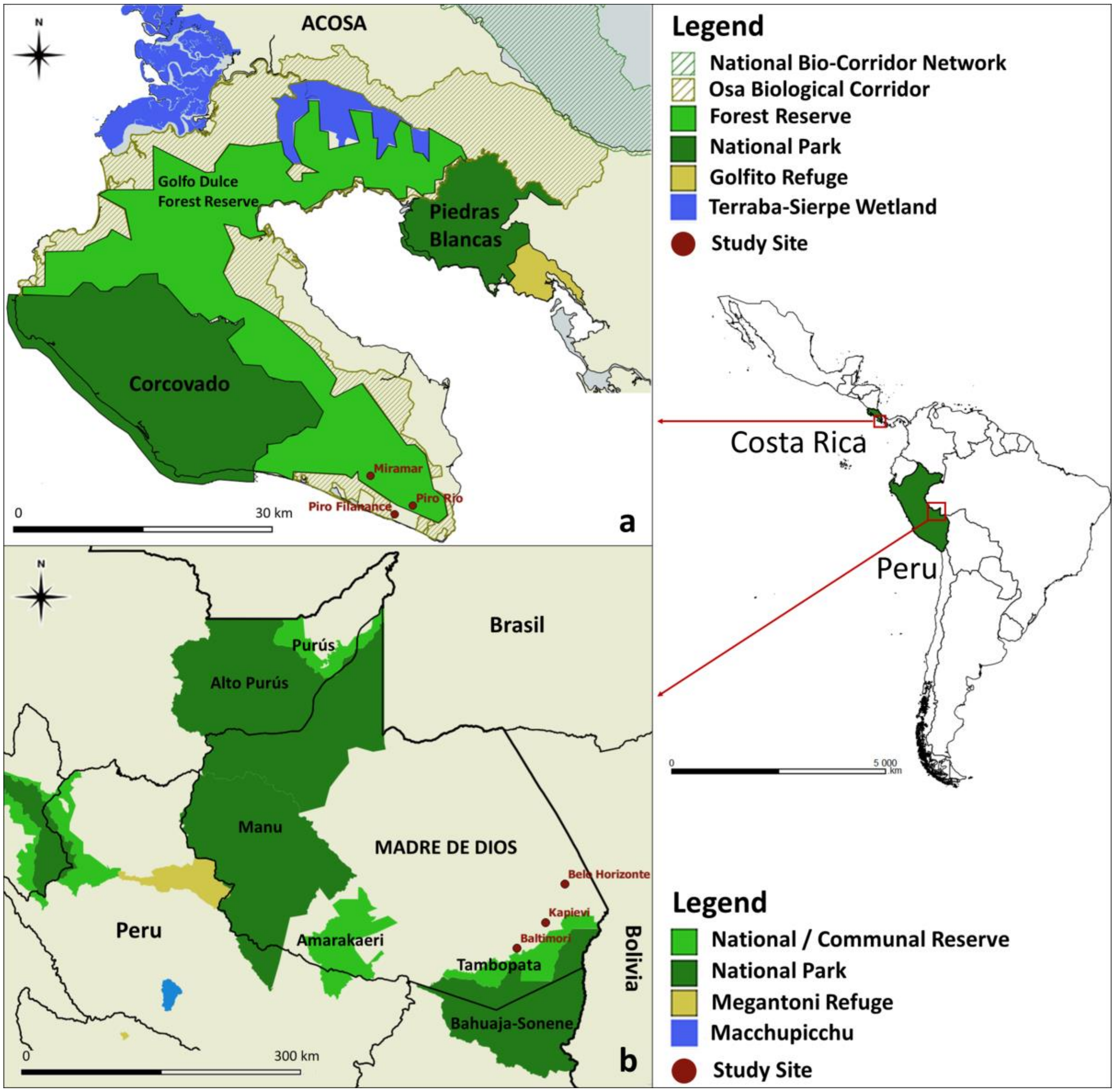

**Figure S1.** Map showing the study sites in (a) Osa Peninsula, Costa Rica (Miramar 8°26’20” N, 83°21’54” W, Piro 8°24’5” N, 83°20’36” W), and (b) Madre de Dios region, Peru (Baltimore 13°23’38” S, 69°44’59” W, Kapievi 13°1’17 S, 69°20’4” W, Belo Horizonte 12°28’38” S, 69°2’59” W). The habitat of the study sites was characterized by secondary tropical rainforests and floodplain wetlands surrounded by humid tropical forest in respectively Costa Rica and Peru. Both areas include various protected areas such as national parks, communal or national forest reserves, and biological corridors. The Osa Peninsula has an average annual precipitation ranging from 2,500 to 6,000 mm, an average annual temperature of 25°C. The annual average rainfall in the Madre de Dios region ranges from 2,000 to 3,500 mm and average temperatures from 21°C to 26°C. The Osa Peninsula hosts six *Vanilla* species: *Vanilla inodora* Schiede, *V. hartii* Rolfe, *V. helleri* A.D. Hawkes, *V. odorata* C. Presl, *V. pompona*, and *V. trigonocarpa* Hoehne (Karremans et al., 2020; Watteyn et al., 2020). The Madre de Dios regions also hosts six species: *Vanilla pompona*, *V. marowynensis* Pulle, *V. karen-christianae* Karremans & P.Lehm., *V. bicolor* Lindl, *V. palmarum* Salzm. ex Lindl., and *V. guianensis* Splitg (Peru) (Householder et al., 2010; Janovec et al., 2013).
