## Appendix B for "Trick or Treat? Pollinator attraction in *Vanilla pompona* (Orchidaceae)"

---

*Bee behavioral comparison analysis*

The observed *V. pompona* floral visitors were assigned to five behavioral categories, following the behavioral hierarchy of bee attraction: (1) approaching, i.e. showing interest in the flower but not necessary landing, (2) landing on the sepals, lateral petals or labellum, (3) landing on the labellum, (4) entering the labellum, and (5) successfully removing the pollen, thus acting as an effective pollinator. The effect of bee genus and time of the day on this ordinal bee behavior classification was modeled using a series of mixed logistic regression models. We sequentially compared each pair of consecutive behavior categories with each other. For example, the first model fits the chance of landing, given a bee approached the flower, as a function of bee genus and time of the day. We accounted for the correlation structure in our data by including random intercepts for flower individual nested within location (Zuur et al., 2009), and the assumption of linearity between the log odds and continuous predictor variable was checked. Due to non-convergence problems with glmmTMB, we applied a Bayesian multilevel model. The models were fit using the R package brms (Bürkner, 2017) with 2 chains, 10,000 iterations and a warm-up of 1000 runs. Using normal priors for intercept and betas, models converged with Rhat values close to 1 (Gelman and Rubin’s diagnostic). Trace plots were used to assess mixing and convergence of the two chains. Parameter estimates of models including only one or both random effects were compared and models were validated via posterior predictive checks, such as the widely applicable information criterion (WAIC; Watanabe, 2010) and the leave-one-out cross-validation (LOO; Gelfand et al., 1992; Vehtari et al., 2017), using the bayesplot R package (Gabry, 2017).

The Bayesian multilevel behavioral comparison analysis showed that most bees approaching the flowers subsequently landed on them (Table S1). Indeed, all of the approaching *Trigona* and respectively 87.7% and 85.7% of the approaching *Eulaema* and *Euglossa* landed on the flower. Closer towards the end of the morning however, the chance that a bee lands on a flower decreases significantly (Figure S2). Additionally, *Euglossa* or *Eulaema* that approach the flowers are predicted to land more on the tepals than on the labellum. When perching on the tepals, both *Euglossa* and *Eulaema* predominantly show a probing behavior (Table S2). The chance that an approaching *Trigona* lands on the labellum is higher compared to *Euglossa* and *Eulaema*, whereby most of the *Trigona* remain on the labellum edge while some enter the labellum tube, all displaying a probing behavior. Nevertheless, *Eulaema* is the only genus that achieves pollen removal when entering the labellum (Figure S3). *Eulaema* or *Euglossa* individuals that enter the labellum, visit the flower for less than five seconds and remain inside the labellum tube for a maximum of two seconds. As they exit, an extended tongue was observed. These rapid visits suggest unsuccessful attempts of nectar foraging. Overall, landing bees display two main reward-approach behaviors on different flower parts: a probing behavior on the tepals (except labellum) or an attempt for nectar search in the labellum (Table S2).

**Table S1.** Output of Bayesian multilevel behavioral comparison models to sequentially compare behavioral categories between floral visitor genera and time of the day. The intercept term is the bee genus *Euglossa* of the categorical variable bee genus. LOOIC = leave-one-out cross-validation, WAIC = Widely Applicable Information Criteria, *SE* = standard error, CI = credibility interval.

| **Behavioral comparison** | **LOOIC** | **WAIC** | **Parameter** | **Estimate** | ***SE*** | **95% CI** |
|---|---|---|---|---|---|---|
| **1. Landing versus approaching** | 74.6 | 64.9 | Intercept | 14.17 | 5.26 | 5.54, 25.97 |
| | | | *Eulaema* | 1.31 | 1.00 | -0.58, 3.32 |
| | | | *Trigona* | 3.42 | 1.49 | 0.64, 6.46 |
| | | | Time of the day | -1.16 | 0.49 | -2.29, -0.39 |
| **2. Landing on labellum versus tepals** | 98.1 | 88.4 | Intercept | -8.77 | 4.91 | -21.60, -2.18 |
| | | | *Eulaema* | 6.49 | 3.86 | 1.87, 16.62 |
| | | | *Trigona* | 16.48 | 7.69 | 7.66, 37.14 |
| | | | Time of the day | -0.01 | 0.21 | -0.43, 0.38 |
| **3. Entering into labellum versus only landing** | 115.1 | 111.1 | Intercept | -5.54 | 2.57 | -11.61, -1.46 |
| | | | *Eulaema* | 3.33 | 1.85 | 0.58, 7.83 |
| | | | *Trigona* | 5.16 | 2.13 | 2.00, 10.36 |
| | | | Time of the day | -0.01 | 0.13 | -0.26, 0.23 |
| **4. Pollen removal versus only entering** | 39.8 | 35.7 | Intercept | -4.70 | 4.57 | -15.82, 2.96 |
| | | | *Eulaema* | 2.65 | 0.88 | 0.97, 4.43 |
| | | | *Trigona* | -0.01 | 0.96 | -1.91, 1.86 |
| | | | Time of the day | 0.12 | 0.47 | -0.69, 1.25 |

**Table S2.** Percentage of landing bees per genus displaying a defined reward approach type on different parts of *Vanilla pompona* flowers (T = tepals; LE = labellum edge; LT = labellum tube), whereby "probing" refers to touching the flower surface with the proboscis and antennae and / or brushing with foretarsi, and "nectar search" refers to entering straight into the labellum chamber, and showing an extended tongue when entering or departing the flower. *N* = total number of landing bees per genus. *SD* = standard deviation.

| **Reward approach** | ***Eulaema* (*N* = 61)** | | | ***Euglossa* (*N* = 23)** | | | ***Trigona* (*N* = 22)** | | | **Visit time (s) (mean ± *SD*)** |
|---|---|---|---|---|---|---|---|---|---|---|
| | T | LE | LT | T | LE | LT | T | LE | LT | |
| **Probing** | 80.3 | 1.6 | - | 95.6 | - | - | 4.5 | 54.6 | 40.9 | 373.5 ± 926.6 |
| **Nectar search** | - | - | 18.1 | - | - | 4.4 | - | - | - | 4.7 ± 2.5 |

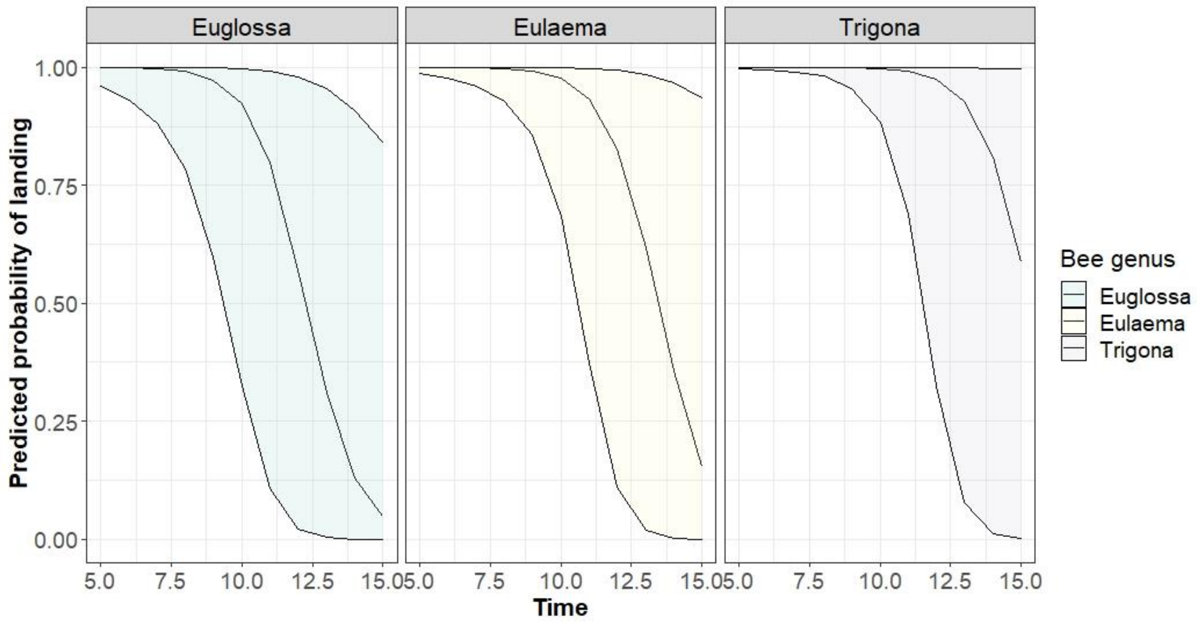


**Figure S2.** Response curve of the behavior landing, shown over time and per bee genus; demonstrating the chance over time of the day that a bee individual belonging to a certain bee genus will land on a *Vanilla pompona* flower.

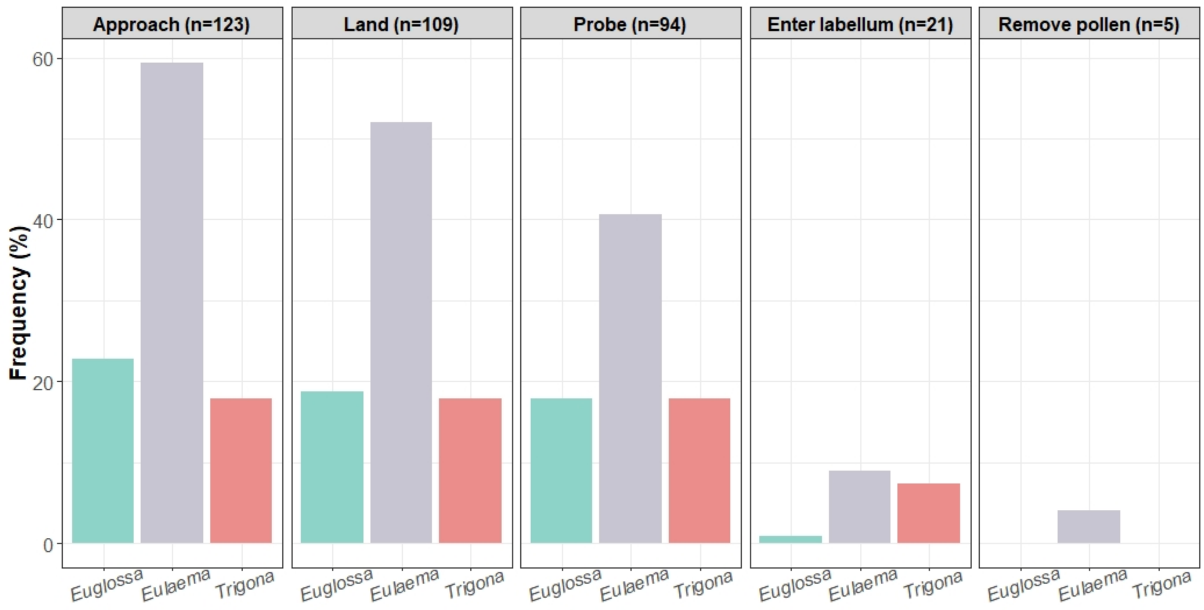


**Figure S3.** Percentage of floral visitors per bee genus that (1) approaches, (2) lands, (3) probes, (4) enters into the labellum, and (5) successfully removes pollen of *V. pompona* flowers.

**References**

Bürkner, P. C. (2017). brms: An R package for Bayesian multilevel models using Stan. *Journal of Statistical Software*, 80(1). doi: 10.18637/jss.v080.i01

Gabry, J. (2017). *Bayesplot: Plotting for Bayesian models*. Retrieved from http://mc-stan.org/bayesplot.

Gelfand, A. E., Dey, D. K., & Chang, H. (1992). Model determination using predictive distributions, with implementation via sampling-based methods. *Bayesian Statistics* 4.

Vehtari, A., Gelman, A., & Gabry, J. (2017). Practical Bayesian model evaluation using leave-one-out cross-validation and WAIC. *Statistics and Computing*, 27(5), 1413-1432.

Watanabe, S., & Opper, M. (2010). Asymptotic equivalence of Bayes cross validation and widely applicable information criterion in singular learning theory. *Journal of machine learning research*, 11(12), 3571-3594.

Zuur, A., Ieno, E. N., Walker, N., Saveliev, A. A, & Smith, G. M. (2009). *Mixed effects models and extensions in ecology with R*, 1st edn. New York: Springer.
