## Appendix C for "Trick or Treat? Pollinator attraction in *Vanilla pompona* (Orchidaceae)"

---

*Nectar*

We applied the rinsing method for small nectar volumes (Morrant et al., 2009; Power et al., 2017) in eight additional flowers. The rinsed water was collected in 2 mL vials and stored at -20°C until High-Performance Liquid Chromatography (HPLC) analysis in an Agilent 1260 Infinity system (Santa Clara, USA) with refractive index detector (G1362A). Three successive rinses (0.5 mL each) of de-ionized UHPLC gradient water were expelled over the labellum. No floral tissue was damaged or removed from the plant to avoid possible leakage of other tissue fluids. Since direct injection of a sample did not detect masses of nectar sugars (sucrose, glucose and fructose), 1 mL of a sample was lyophilized using 1-propanol HPLC grade and evaporated with nitrogen. This was then diluted in 30 µL mobile phase (ultrapure water type 1, Milli-Q®, Millipore Corporation, Burlington, USA) to concentrate the sugars 33 times more to quantify them. Sugar separation was achieved by injecting 9 to 15 µL of this solution into a Hi-Plex Ca2+ column (300 × 7.7 mm, 8 µm, Agilent, Santa Clara, USA) at 45 °C, in isocratic mode with a flow of 0.5 mL/min. A 25 min chromatographic run achieved separation and the limits of detection (LOD) was determined by the signal-to-noise ratio (S/N ratio = 3) (Figure S4).

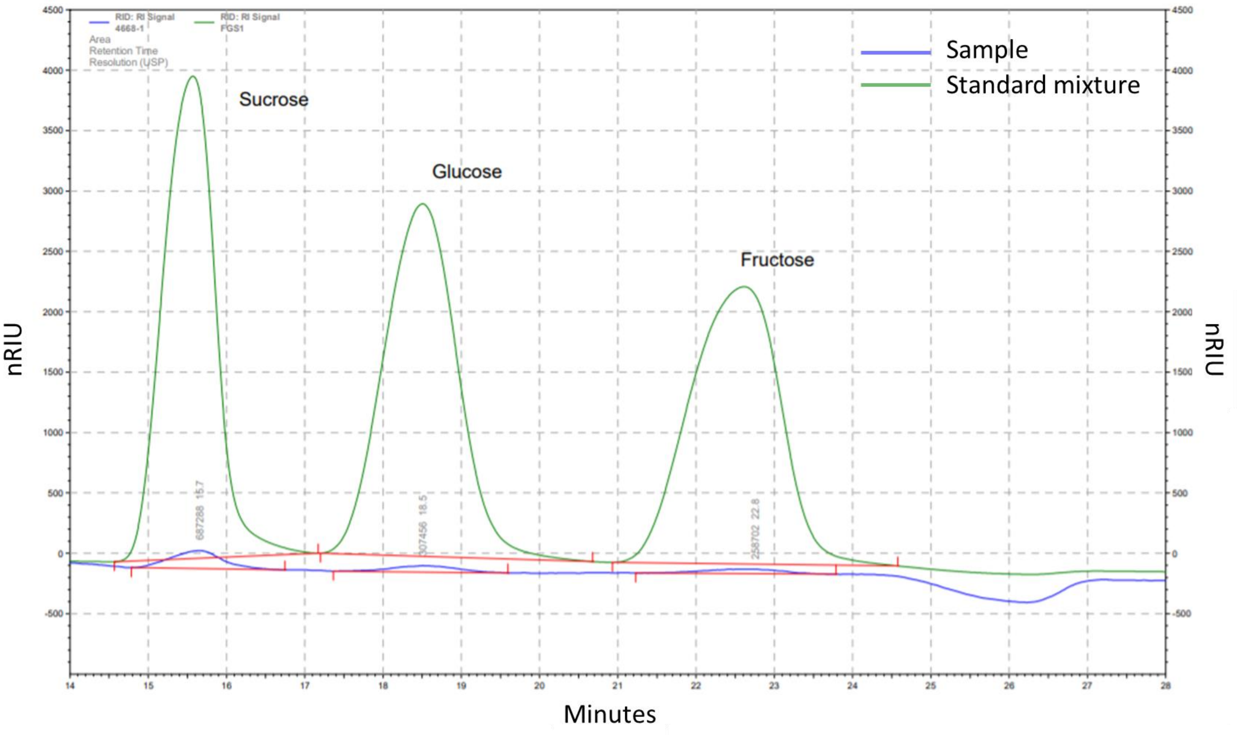


**Figure S4.** HPLC-chromatogram of a *V. pompona* sample extracted from the labellum via three successive rinses (0.5 mL each) of de-ionized UHPLC gradient water. The green line represents a standard mixture of three sugars (sucrose, glucose, fructose). The blue line shows the sugar content of the sample. The limits of detection (LOD) was determined by the signal to noise (S/N) ratio method and estimated as the minimum concentration providing a S/N ratio of 3:1.

*Floral volatiles*

Floral fragrances of 12 flowers were collected by static headspace sampling (Tholl et al., 2006) during the flowering period of 2020 in two Costa Rican populations. The sampling was conducted between 7:00 am and 11:00 am, corresponding to the activity peak of the potential pollinators (Dodson et al., 1969; Ackerman, 1983b; Armbruster et al., 2000; Gostinski et al., 2016), thus expected floral scent productivity peaks (Whitten, 1985; Hills & Williams, 1990). The single-use scent traps were constructed using clear glass tubing (2.4 mm ID, 3.5 cm length). They were plugged at both ends with glass wool, filled with 20 mg of bulk carbon and 20 mg of Tenax GC (Williams & Whitten, 1983; Whitten, 1985; Raguso & Pellmyr, 1998), and conditioned by passing 5 mL of hexane. During each sampling, one inflorescence was bagged with a plastic scentless oven bag (Reynolds Kitchens) closed at the top for 30 min. The scent traps were then connected with the tenax-side to the bag and the carbon-side to a battery-operated vacuum pump via Tygon tubing (ID 3.3 mm). For three hours, air was continuously extracted from the bag through the scent trap and the flower rate was adjusted to 200 mL/min using a 9 V battery. The scent traps were eluted with 200 μL of hexane into conical inserts held in 2 mL auto-sampler vials (Agilent Technologies), stored at -20°C until analysis. Control samples were collected simultaneously and in the same manner from bags filled with ambient air.

The scent traps were analyzed by gas chromatography-mass spectrometry (GC-MS) analysis, using an Agilent 7890B GC fitted with a 30 m x 0.25 mm x 0.25 µm HP-5 Ultra Inert column coupled to an Agilent 5977A mass spectrometer (Agilent Technologies). We used the same methodology for analysis, data processing, compound identification, and compound attractiveness as described in Hetherington-Rauth & Ramirez (2016). Presumed contaminant compounds were removed when present in both control and sample to obtain the final set of floral volatile compounds. We compiled information on the attractiveness of each of the final compounds from the literature, including field behavioral assays (Dodson et al., 1969; Williams & Dodson, 1972; Williams & Whitten, 1983; Ackerman, 1989; Hetherington-Rauth & Ramirez, 2016). The compound peak areas of the final set of floral volatiles were converted to relative frequencies and averaged per compound among the samples to reveal potential patterns within the floral bouquet (Figure S5).

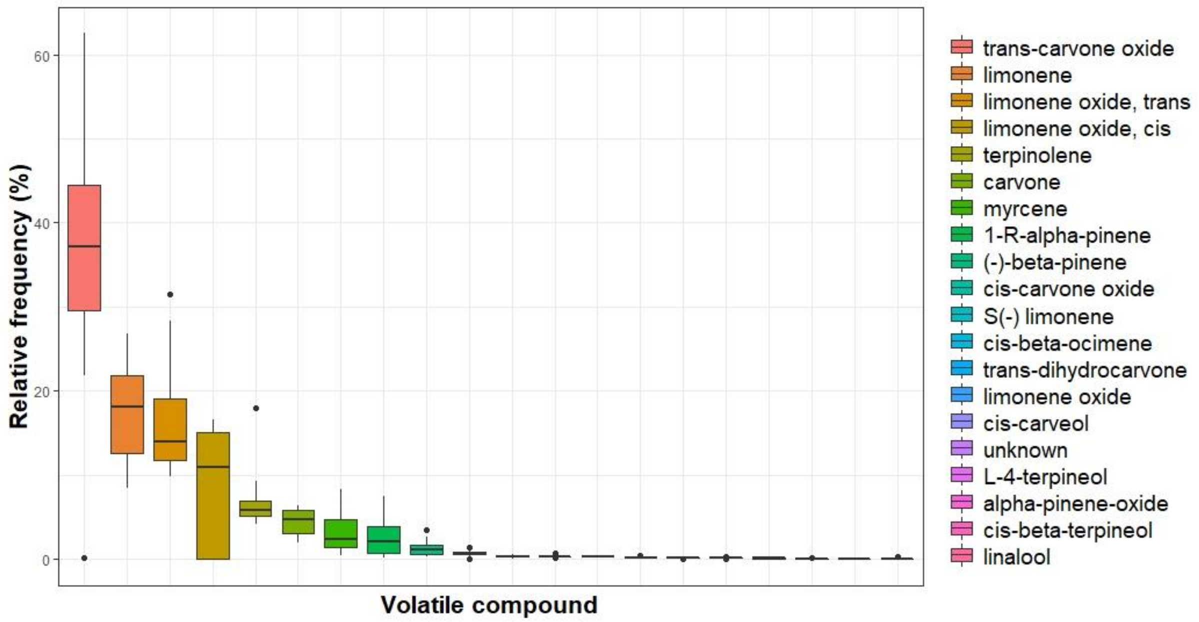


**Figure S5.** Floral volatile compound composition of *Vanilla pompona* flowers, calculated by averaging the relative frequency of each individual compound among the twelve samples.

**References**


Morant, D. S., Schumann, R., & Petit, S. (2009). Field methods for samping and storing nectar from flowers with low nectar volumes. *Annals of Botany*, 103(3), 533-542.

Power, E. F., Stabler, D., Borland, A. M., Barnes, J., & Wright, G. A. (2018). Analysis of nectar from low-volume flowers: A comparison of collection methods for free amino acids. *Methods in ecology and evolution*, 9(2), 734-743.

Raguso, R. A., & Pellmyr, O. (1998). Dynamic headspace analysis of floral volatiles: a comparison of methods. Oikos, 81(2), 238-254.

Tholl, D., Boland, W., Hansel, A., Loreto, F., Röse, U. S. R., & Schnitzler, J. P. (2006). Practical approaches to plant volatiles analysis. *The Plant Journal*, 45(4), 540-560.

Dodson, C. H., Dressler, R. L., Hills, H. G., Adams, R. M., & Williams, N. H. (1969). Biologically active compounds in orchid fragrances. *Science,* 164 (3885), 1243-1249.

Ackerman, J. D. (1983b). Diversity and seasonality of male euglossine bees (Hymenoptera: Apidae) in Central Panama. *Ecology*, 64(2), 274-283.

Armbruster, W. S., Fenster, C., & Dudash, M. (2000). Pollination "principles" revisited: specialization, pollination syndromes, and the evolution of flowers. *The Scandanavian Association for Pollination Ecology*, 39, 179-200.

Gostinski, L. F., Carvalho, G. C. A., Rego, M. M. C., & Albuquerque, P. M. C. (2016). Species richness and activity patterns of bees (Hymenoptera: Apidae) in the restinga area of Lençóis Maranhenses National Park, Barreirinhas, Maranhão, Brazil. *Revista Brasileira de Entomologia*, 60(4), 319-327.

Hills, G. H., & Williams, N. H. (1990). Fragrance cycle of *Clowesia rosea*. *Orquidea*, 12, 119-132.

Williams, P. H., & Whitten, W. M. (1983). Orchid floral fragrances and male euglosssine bees: methods and advances in the last sesquidecade. *Biological Bulletin*, 164(3), 355-395.

Whitten, W. M. (1985). *Variation in floral fragrances and pollinators in the Gongora quinquenervis complex (Orchidaceae) in Central Panama*. PhD Thesis, Florida University, USA.

Williams, N. H., & Dodson, C. H. (1972). Selective attraction of male euglossine bees to orchid floral fragrances and its importance in long-distance pollen flow. *Evolution*, 26(1), 84-95.

Ackerman, J. D. (1989). Geographic and seasonal variation in fragrance choices and preferences of male euglossine bees. *Biotropica*, 21(4), 340-347.

Hetherington-Rauth, M. C., & Ramirez, S. R. (2016). Evolution and diversity of floral scent chemistry in the euglossine bee-pollinated orchid genus *Gongora*. *Annals of Botany*, 118(1), 135-148.
